# Usefulness of docking and molecular dynamics in selecting tumor neoantigens to design personalized cancer vaccines: *a proof of concept*

**DOI:** 10.1101/2022.12.22.521559

**Authors:** Diego Amaya-Ramirez, Laura Camila Martinez-Enriquez, Carlos Parra-López

## Abstract

Personalized cancer vaccines are presented as a new and promising treatment alternative for cancer, especially in those cases where effective treatments do not yet exist. However, multiple challenges remain to be resolved so that this type of immunotherapy can be used in the clinical setting. Among these, the effective identification of immunogenic peptides stands out, since the *in-silico* tools currently used generate a significant portion of false positives. This is where molecular simulation techniques can play an important role when it comes to refining the results produced by these tools. In the present work, we explore the use of molecular simulation techniques such as docking and molecular dynamics to study the relationship between stability of peptide-HLA complexes and their immunogenicity using two HLA-A2-restricted neoantigens that have already been evaluated *in vitro*. The results obtained agreed with the *in vitro* immunogenicity of the immunogenic neoantigen ASTN1 the only one that remains bound at both ends to the HLA-A2 molecule. Additionally, molecular dynamics indicates that position 1 of the peptide has a more important role in stabilizing the N-terminal part than previously assumed. Likewise, the results suggest that the mutations may have a “delocalized” effect on the peptide-HLA interaction, that is, they may modulate the intensity of the interactions of other amino acids in the peptide. These results highlight the suitability of this type of *in silico* strategy to identify peptides that form stable complexes with HLA proteins that are highly immunogenic for CD8+ T cells.

## Introduction

Personalized neoantigen-based vaccines have proven to be a useful tool for immunotherapy of aggressive tumors, such as metastatic melanoma, glioblastoma, and non-small cell lung cancer (1–6). This type of vaccine promotes a highly specific response of T lymphocytes that did not undergo the tolerance induction process in the thymus because they recognize neoantigens encoded in somatic mutations of the tumor (7–9).

Currently, tumor neoantigens are predicted using *in silico* strategies, which compare the tumor DNA sequence with that of healthy patient tissue to identify tumor-specific somatic mutations. Once the mutations have been identified, the neoantigens are predicted by using trained algorithms to estimate the probability that they will be processed and presented in the context of Major Histocompatibility Complex (MHC) molecules, also called Human Leukocyte Antigen (HLA), of the patient (10–15). Once the identity of the neoantigens has been established, the sequence of the tumor transcriptome allows the expression levels of the predicted neoantigens to be verified. Other parameters incorporated in bioinformatic tools for the identification of immunogenic neoantigens are the relative affinity of the MHC molecule neoantigen complex (IC50) (16–18), and the half-life of the binding of the MHC-neoantigen complex (Stability) (19).

These *in silico* strategies allow to prioritize potentially immunogenic neoantigens. However, despite the promising results of some studies on the design and clinical response of tumors to personalized vaccines, clinical studies have shown a limited *in vivo* immunogenicity of the selected neoantigens (20), due probably to a limited capacity of *in silico* methods to identify immunogenic epitopes *in vivo*. Despite the progress of *in silico* prediction tools, they still have a high rate of false positives (low specificity) (21, 22), which is mainly due to two factors: 1) the dataset used to train these tools is usually based on information generated from a limited number of HLA alleles, and, they do not take into account results of *in vitro* or *in vivo* evaluation of neoantigens; and 2) Predictive tools rely on sequence data only and therefore do not satisfactorily incorporate molecular aspects of epitope processing and presentation by antigen-presenting cells, such as stability of the MHC-peptide complex and recognition of the MHC-peptide complex by the T cell Receptor (TCR) on T cells, that are important factors determining the immunogenicity of antigens (23).Therefore, it is necessary to search for new tools to improve the selection of peptides with immunogenic potential *in vivo*.

Docking and molecular dynamics are computational tools that allow the understanding of non-covalent receptor-ligand interactions at atomic level useful in rational drug design (24), and in the study of the interaction of HLA-peptide-complex with the TCR molecules (25–29). The use of these tools in the selection of immunogenic tumor neoantigens have not been explored. Docking and molecular dynamics might make it possible to generate additional information on the interaction of HLA molecules with neoantigens, and perhaps, to a better selection of neoantigens efficiently presented by MHC molecules and recognized by T cells *in vivo*.

In this work, molecular docking and molecular dynamics were used to discriminate molecular interactions among HLA-A*02:01 molecules and two neoantigens one immunogenic and one non-immunogenic for T cells. Based on the study by Strønen et al., (30) we choose two melanoma neoantigens that according to predictive algorithms bind to HLA-A*02:01 molecules with high affinity and whose data on immunogenicity were available. To elucidate the structural properties of a neoantigen in complex with HLA-A*02:01 leading the expansion of human cytotoxic CD8+ T-cell that efficiently recognize and destroy melanoma cells, we analyzed both complexes through the aforementioned techniques and found remarkably different structural features on each complex.

## METHODOLOGY

### Neoantigen selection

Based on the study published by Strønen et al. (30), 21 neoantigens and their wild-type counterparts were selected (see Table 1). The peptides had the following characteristics: (i) the amino acid sequence of the mutated peptide includes a single amino acid change compared to the wild-type sequence, (ii) the affinity score of the mutant peptide was <1000nM (according to NetMHC 3.2) and (iii) both, the stability of the peptide-HLA-A*02:01 complexes were measured and (iii) the immunogenicity for CD8+ T cells were assessed experimentally.

**Table 1:**
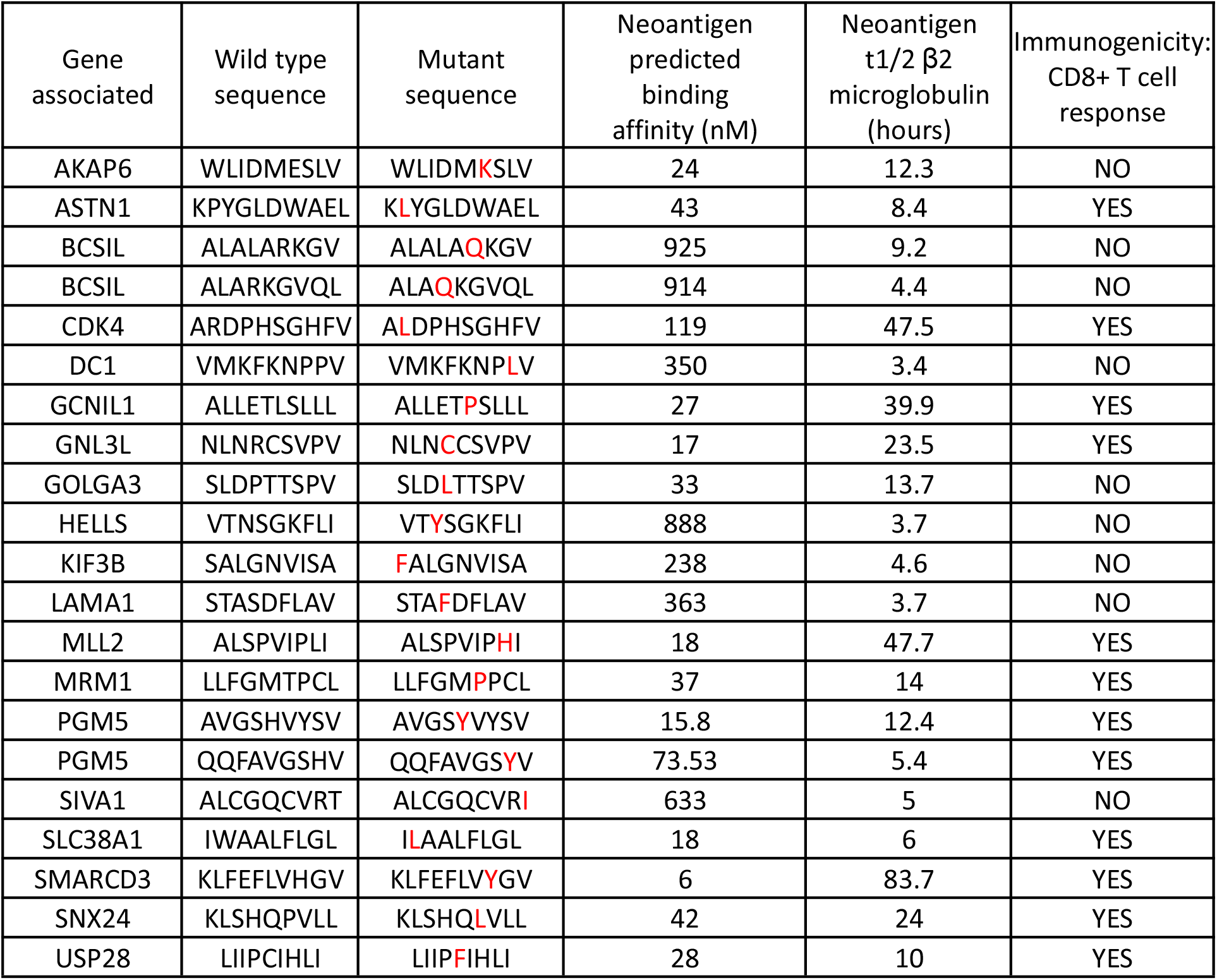
Peptide sequences of the wild type and mutant neoantigen selected for the present study. Highlighted in red are the positions where the mutations of the neoantigens are located with respect to their wild-type counterparts.

The peptide-HLA-A*02:01 complexes of these 42 peptides were modeled by molecular docking and then analyzed with LigPlot+ (31) to select two neoantigens and their wild-type counterparts for molecular dynamic simulation. The selection was based on the number of total interactions between the peptide and HLA, and the immunogenicity reported by Stronen. Considering these criteria, the wild-type and neoantigen associated with the gene ASTN1 (10 amino acids in length) and AKAP6 (9 amino acids in length) were chosen as immunogenic and non-immunogenic neoantigens respectively.

### Evaluation of peptides through sequence based in silico tools

The NetMHC 4.0 tool for HLA-A*02:01 binding affinity assessment (32), NetMHCstabpan 1.0 to predict MHC-I binding stability (19), NetCTL 1.2 and NetTepi 1.0 to determine proteasome processing and TAP transport and HLA binding (33, 34), were all used to predict immunogenicity for CD8+ T cells of the four peptide sequences listed in Table 3.

**Table 2.** Number of total interaction and hydrophobic interaction.

| Peptide | Sequence | # Total interactions | # Hydrophobic interactions | *In vitro* immunogenicity |
| --- | --- | --- | --- | --- |
| AKAP6 | WLIDMKSLV | 29 | 18 | NO |
| DC1 | VMKFKNPLV | 28 | 18 | NO |
| ASTN1 | KLYGLDWAEL | 26 | 16 | YES |
| GCN1L1 | ALLETPSLLL | 25 | 14 | YES |
| GNL3L | NLNCCSVPV | 25 | 14 | YES |
| HELLS | VTYSGKFLI | 24 | 15 | YES |
| SMARCD3 | KLFEFLVYGV | 24 | 13 | YES |
| USP28 | LIIPFIHLI | 24 | 15 | YES |
| GOLGA3 | SLDLTTSPV | 23 | 13 | NO |
| MRM1 | LLFGMPPCL | 23 | 15 | YES |
| PGM5 | QQFAVGSYV | 23 | 11 | YES |
| MLL2 | ALSPVIPHI | 22 | 13 | YES |
| SIVA1 | ALCGQCVRI | 21 | 11 | NO |
| SLC38A1 | ILAALFLGL | 21 | 13 | YES |
| BCSIL | ALAQKGVQL | 20 | 11 | NO |
| CDK4 | ALDPHSGHFV | 19 | 10 | YES |
| LAMA1 | STAFDFLAV | 19 | 9 | NO |
| SNX24 | KLSHQLVLL | 19 | 10 | YES |
| BCSIL | ALALAQKGV | 18 | 11 | NO |
| KIF3B | FALGNVISA | 18 | 9 | NO |
| PGM5 | AVGSYVYSV | 16 | 8 | YES |

**Table 3.**
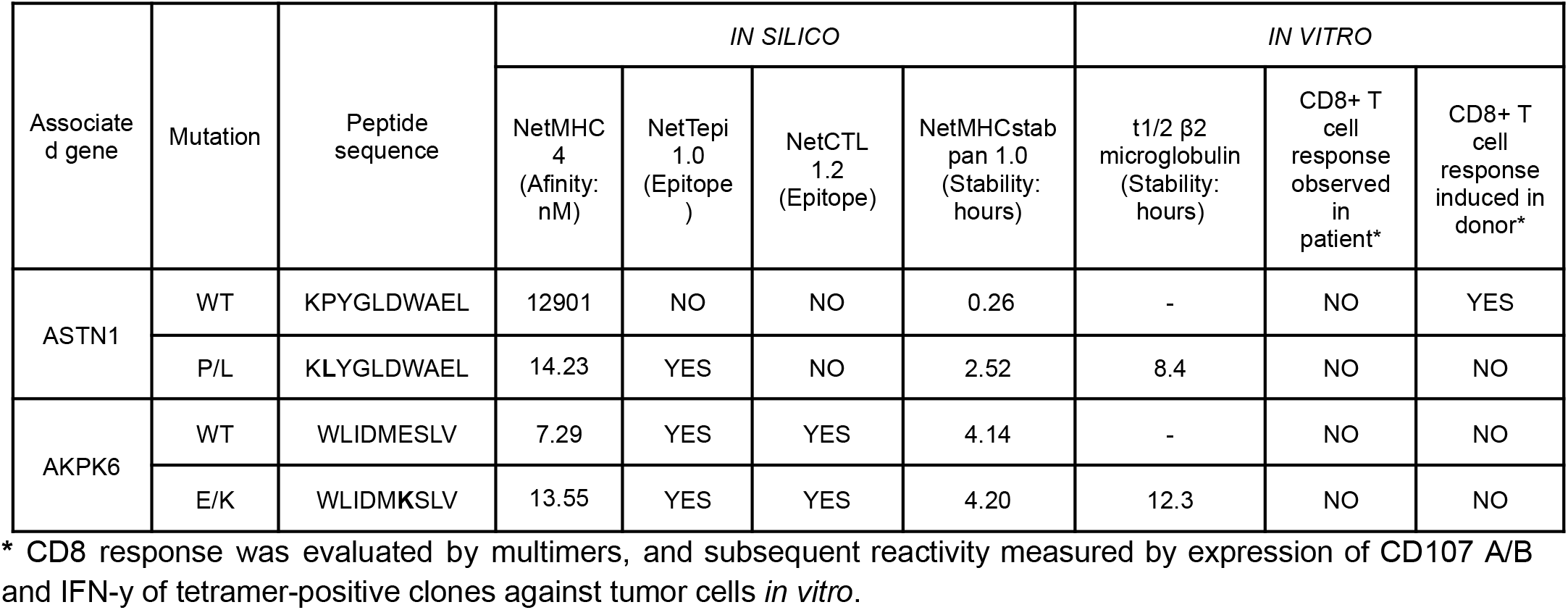
Results of analyses of four peptides assessed by using the different tools traditionally used for the identification of neoantigens.

| Associate d gene | Mutation | Peptide sequence | IN SILICO |  |  |  | IN VITRO |  |  |
| --- | --- | --- | --- | --- | --- | --- | --- | --- | --- |
| | | | NetMHC 4<br>(Afinity: nM) | NetTepi 1.0<br>(Epitope ) | NetCTL 1.2<br>(Epitope) | NetMHCstab pan 1.0<br>(Stability: hours) | t1/2 β2 microglobulin<br>(Stability: hours) | CD8+ T cell response observed in patient* | CD8+ T cell response induced in donor* |
| ASTN1 | WT | KPYGLDWAEL | 12901 | NO | NO | 0.26 | - | NO | YES |
|  | P/L | KLYGLDWAEL | 14.23 | YES | NO | 2.52 | 8.4 | NO | NO |
| AKPK6 | WT | WLIDMESLV | 7.29 | YES | YES | 4.14 | - | NO | NO |
|  | E/K | WLIDMKSLV | 13.55 | YES | YES | 4.20 | 12.3 | NO | NO |

\* CD8 response was evaluated by multimers, and subsequent reactivity measured by expression of CD107 A/B and IFN- $\gamma$ of tetramer-positive clones against tumor cells *in vitro*.

### Molecular docking

To generate the peptide-HLA complex model for each case, molecular docking was performed. Whereas to model the peptide pair derived from ASTN1 (wild-type and mutant neoantigen), the structure 5C0G from the Protein Data Bank (PDB) (35) was used as a reference structure, for the pair of peptides derived from AKAP6, the 5NMH structure was used. The FlexPepDock Ab-Initio protocol (36) integrated in the Rosetta package (37) was implemented for docking, where 50,000 poses were calculated for each peptide. According to the Rosetta scoring function, the best pose generated for each peptide was selected. The number of interactions (Hydrogen bonds and hydrophobic interactions) of the peptide-MHC complex were evaluated through Ligplot+ v2.2.

### Molecular dynamics simulation

Subsequently, a molecular dynamics simulation was performed for each peptide using the VMD (38) and NAMD (39) tools. For the generation of the topology of the system, the VMD AutoPSF tool (38) was used together with the CHARMM36 force field (40), the system was solvated using the TIP3P explicit solvent model, the size of the water box was 67×70×70 A^3^, the system was neutralized using the VMD AutoIonize tool (38), the Particle Mesh Ewald (PME) method was used to calculate the electrostatic energy with a distance truncation of 11 Å. The simulation was carried out under NPT conditions, that is, constant pressure (1 atm) and temperature (310 K) and was composed of 3 stages: (i) minimization, for which the system was brought into room temperature (310K), (ii) system stabilization for ~15ns and (iii) the simulation itself with a duration of 295ns. Lastly, the VMD (38) and Pycontact (41) tools were used to analyze, both, the peptide-protein interactions and to assess the complex stability.

## RESULTS

Initially, molecular docking was carried out to model the peptide-MHC complex (p-MHC) of the 21 neoantigens listed in Table 1 and their wild-type counterparts. The best pose of each of the peptides on HLA-A*02:01 was analyzed using the Ligplot+ tool to determine the number of hydrophobic interactions and hydrogen bonds in the p-MHC complex for both the mutant version (Table 2) as the wild version (Table S1). Overall, the total interactions range from 16-29 without observing a pattern that allows us to correlate immunogenicity with the number of interactions. We proceeded to select two neoantigens, one immunogenic and the other non-immunogenic, to perform the simulation of the p-MHC complex through molecular dynamics to analyze the peptide-protein interactions and the stability of the complex over time. Therefore, based on the number of interactions by molecular docking and the immunogenicity reported by Strønen, the AKAP6 peptide with 29 total interactions was selected as non-immunogenic neoantigen, and ASTN1 with 26 total interactions as immunogenic neoantigen to continue with the molecular simulations.

### Limitation of the predictive tools based on sequence

The AKAP6 neoantigen was generated by substituting glutamic acid for a lysine (E/K) at position 6 (P6). The predictive analysis of the AKPK6 neoantigen sequence (IC50 and complex stability values) suggested that both the wild type and the mutant sequences should be more immunogenic than the ASTN1 neoantigen for CD8+ T-cells; however, the *in-vitro* evaluation carried out by Strønen, proved that neither the wild type nor the mutant sequence of AKAP6 were immunogenic. On the other hand, the ASTN1 neoantigen is generated by a change of proline by a leucine (P/L) at position 2 (P2). Whereas the NetCTL tool (that predicts cleavage by the proteasome and efficiency of transport by TAP) revealed that ASTN1 does not meet characteristics of an immunogenic sequence, NetMHC 4.0 and NetMHCstabpan 1.0 tools revealed that ASTN1 neoantigen has a higher affinity for the HLA-A *0201 molecule and forms stable MHC-peptide complexes predicting a more immunogenic sequence than that formed by the wild-type sequence. This has been proved experimentally by Strønen, since the *in vitro* evaluation of this neoantigen clearly demonstrated that this neoantigen is highly immunogenic for CD8-T cells (Table 3). Altogether, our results suggest that predictive algorithms provide conflicting results that are hard to conceal with epitope immunogenicity and argue for the need to have other techniques to improve the prediction of immunogenic epitopes.

### Molecular simulations

From the structures derived from molecular docking for AKAP6 and ASTN1, both for the wild-type versions and for the neoantigens, a peptide-HLA binding is observed with a conventional orientation with the side chains of the P2 and P9/P10 residues arranged in the pockets of HLA-A2 (Figure 1A and B). For AKAP6: P2 is Leucine and P9 is Valine; for ASTN1: P2 is Proline/Leucine (WT/Neo), and P10 is Leucine. The exposed side chains for AKAP6 are P1, P5, and P8, and for ASTN1 are P1, P5, P6, P8, and P9, all of which project away from the HLA-A2 binding pockets forming a surface with the potential to interact with HLA-A2.

**Figure 1.**
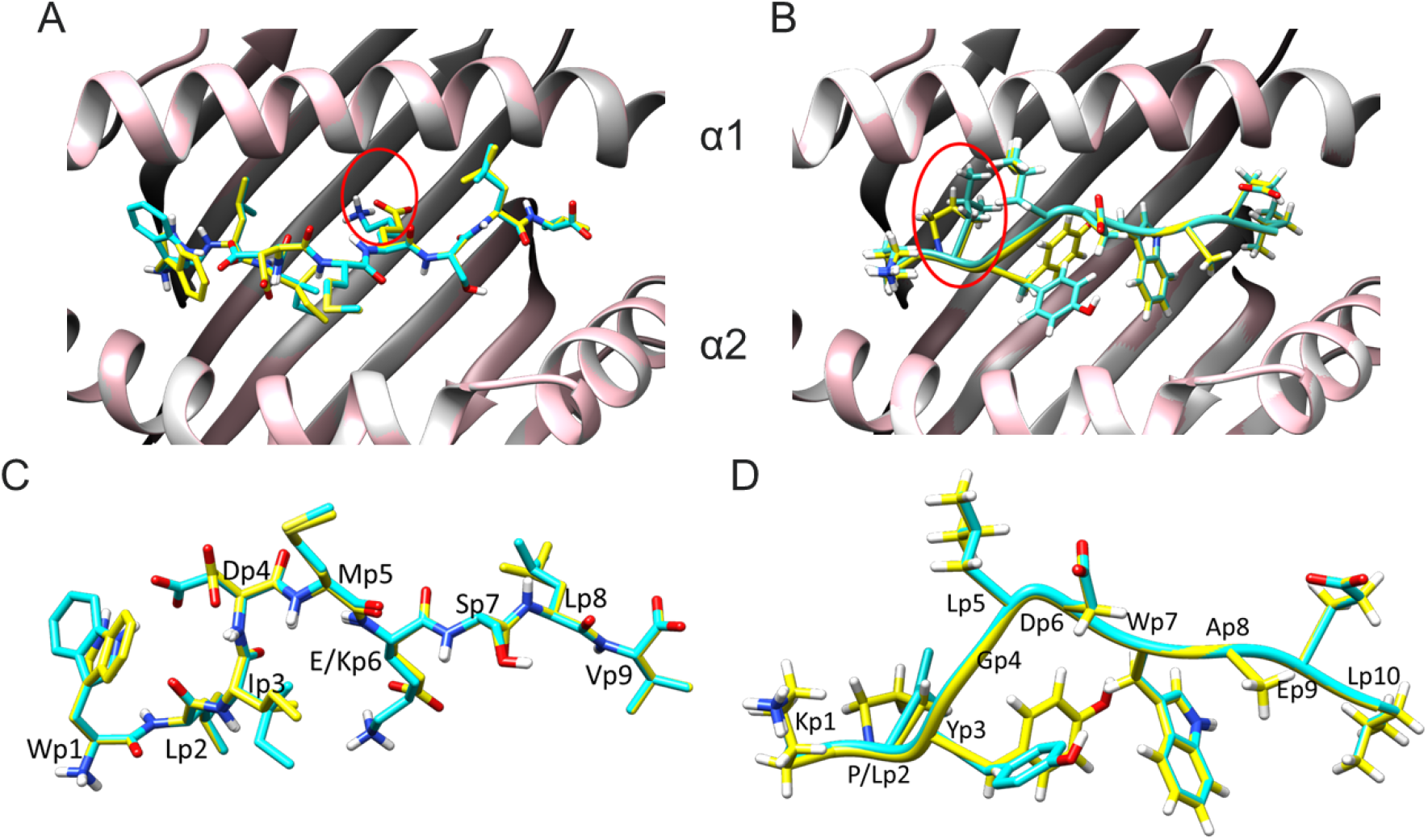
Conformers of wild-type and mutant peptides for AKPK6 and ASTN1 bound to HLA-A*02:01. (A) Top view of AKAP6wt-HLA-A2 and AKAP6 neo-A2 complexes. (B) Top view of ASTN1wt-HLA-A2 and ASTN1 neo-A2 complexes. The wild version is in yellow, and the mutant is in cyan. The point mutation is marked in a red circle. The HLA-A2 backbone is in white (WT-HLA-A2) or pink (neo-HLA-A2). C. Side view of overlapping AKAP6 mutant and wild-type peptides. (D) Side view of overlapping wild-type and mutant ASTN1 peptides. Carbon atoms are in yellow (WT) or in cyan (Neo); nitrogen atoms are in blue; oxygen atoms are in red.

In both cases, the peptide-MHC complexes show structural deviations between the wild-type version and the neoantigen (Figures 1C and D). In particular, the wild type and mutant peptides of AKAP6 overlap in all the side chains except for P1, P3, and P6, being the last ones where the mutation occurs. For its part, in ASTN1, only P2 and P3 change, being the first where the mutation occurs. Therefore, the structural differences between the peptide-MHC complexes for the wild-type and the mutant version in both neoantigens appear not to be restricted solely to the mutation site.

The molecular dynamics simulations showed very different structural attributes of both neoantigens bound to the HLA-A*0201 molecule. On one hand, for AKAP6, both the wild-type (link to visualize online the simulation: https://mmb.irbbarcelona.org/3dRS/shared/61b278eec0a8d2.72605506, <u>corresponding files are also available in supplemental materials</u>) and the mutant version ( link to visualize online the simulation: https://mmb.irbbarcelona.org/3dRS/shared/61b27a3cdfdef2.66763559, <u>corresponding files are also available in supplemental materials</u>) binds to the HLA-A*0201 molecule, however, by the end of the simulation, the C-terminal end of both peptides dissociate from the peptide binding groove (PBG) of the HLA-A*0201 molecule, remaining anchored only by the N-terminal (P1 to P5). In the case of ASTN1, the simulations showed that the wild-type version of the peptide (link to visualize online the simulation: https://mmb.irbbarcelona.org/3dRS/shared/61b27dac052904.99817841, <u>corresponding files are also available in supplemental materials</u>) detaches from the N-terminal end and remains anchored to the MHC molecule through the C-terminal end (P7 to P10). In contrast, the neoantigen sequence of ASTN1 remains anchored at both ends of the MHCI throughout the simulation time, meaning that the amino acid change in P2 allows ASTN1 neoantigen to form the most stable peptide-HLA-A*0201 complex of the four MHCI-peptide complexes analyzed (link to visualize online the simulation: https://mmb.irbbarcelona.org/3dRS/shared/61b27c62dc9300.82271714, <u>corresponding files are also available in supplemental materials</u>).

### Atomic interactions

Atomic interactions between peptide and HLA-A* 0201 were analyzed using the Pycontact tool (41). Pycontact is a bioinformatics tool that allows to identify and characterize noncovalent interactions between molecules in a molecular dynamics simulation. Particularly, it allows calculating the intensity of interaction through a magnitude called “contact score.” That is, the stronger the interaction between two atoms/residues, the higher the “contact score.” This analysis focused on stable interactions over time, that is, those interactions with a median “contact score” greater than zero. At first, the number and type of stable interactions were analyzed, with the predominant types of interactions being hydrophobic and hydrogen bonds (Figure 2). Regarding the types of interactions considered here, it is worth clarifying that the “other” category corresponds to those interactions that do not strictly meet the classification thresholds of any of the other categories.

**Figure 2.**
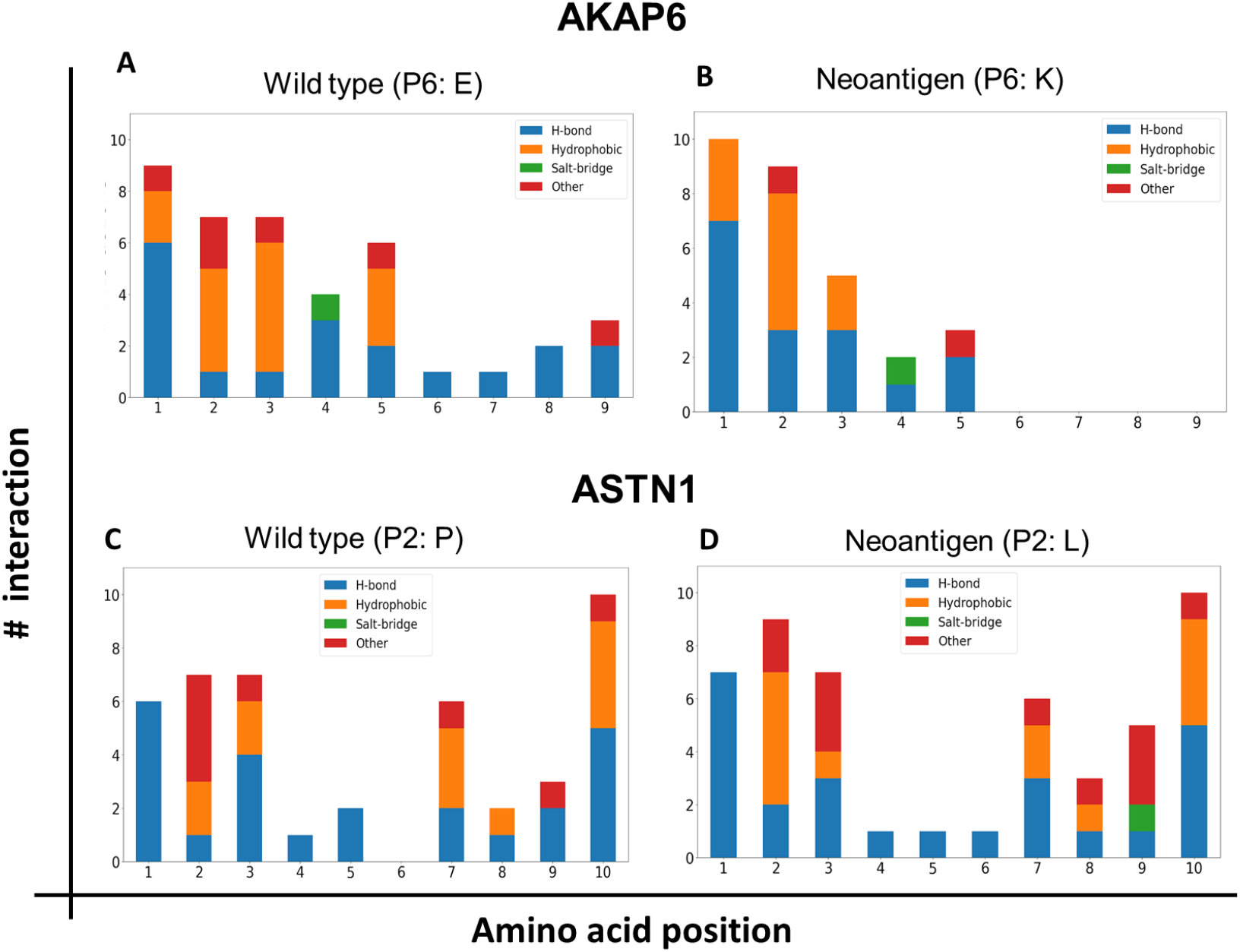
Number and type of interactions between each of the amino acids of the antigens and HLA-A*02:01. Molecular dynamics simulations were performed for the wild type and the mutant version for both neoantigens, being able to analyze the number and type of noncovalent interactions between the amino acids of the epitopes and HLA-A2 using the Pycontact tool. A and B bar graphs for the non-immunogenic neoantigen AKAP6. C and D bar graphs for the immunogenic neoantigen ASTN1. In blue hydrogen bonds, in orange hydrophobic interactions, in green salt bridges and, in red other types of interactions.

In the case of AKAP6, even though both the wild-type peptide and the neoantigen detach from the C-terminal end (more precisely, positions P6 to P9), the mutated peptide presents a reduction in the number of interactions in the C-terminal part generating faster release compared to the wild-type version (Figure 2A). In the case of ASTN1, the mutation in this neoantigen generates an increase in the number of stable interactions with HLA not only at the position where the amino acid change occurred (P2) but also at other positions (notably at the P9). This increase in interactions favors the stability of the complex, as evidenced in the molecular simulation since it is this peptide that remains anchored at both ends, contrary to the wild type that is released from the N-terminal end.

On the other hand, the variability of the types of interaction between neoantigens and their wild-type is striking. This illustrates the complexity of the dynamics of protein-peptide interactions and helps to understand why current tools trained primarily on sequence information are unable to accurately predict the stability of peptide-HLA complexes.

### Intensity of interactions (contact score)

The intensity of interaction between atoms/residues is of great interest when it comes to characterizing the stability of a complex. In AKAP6, a significant increase in the average intensity of interaction in P3 and P4 between the wild type and the mutated version is observed (Figure 3). However, this change is irrelevant in the global stabilization of the peptide since, in both cases, the peptide is released from the C-terminal end.

**Figure 3.**
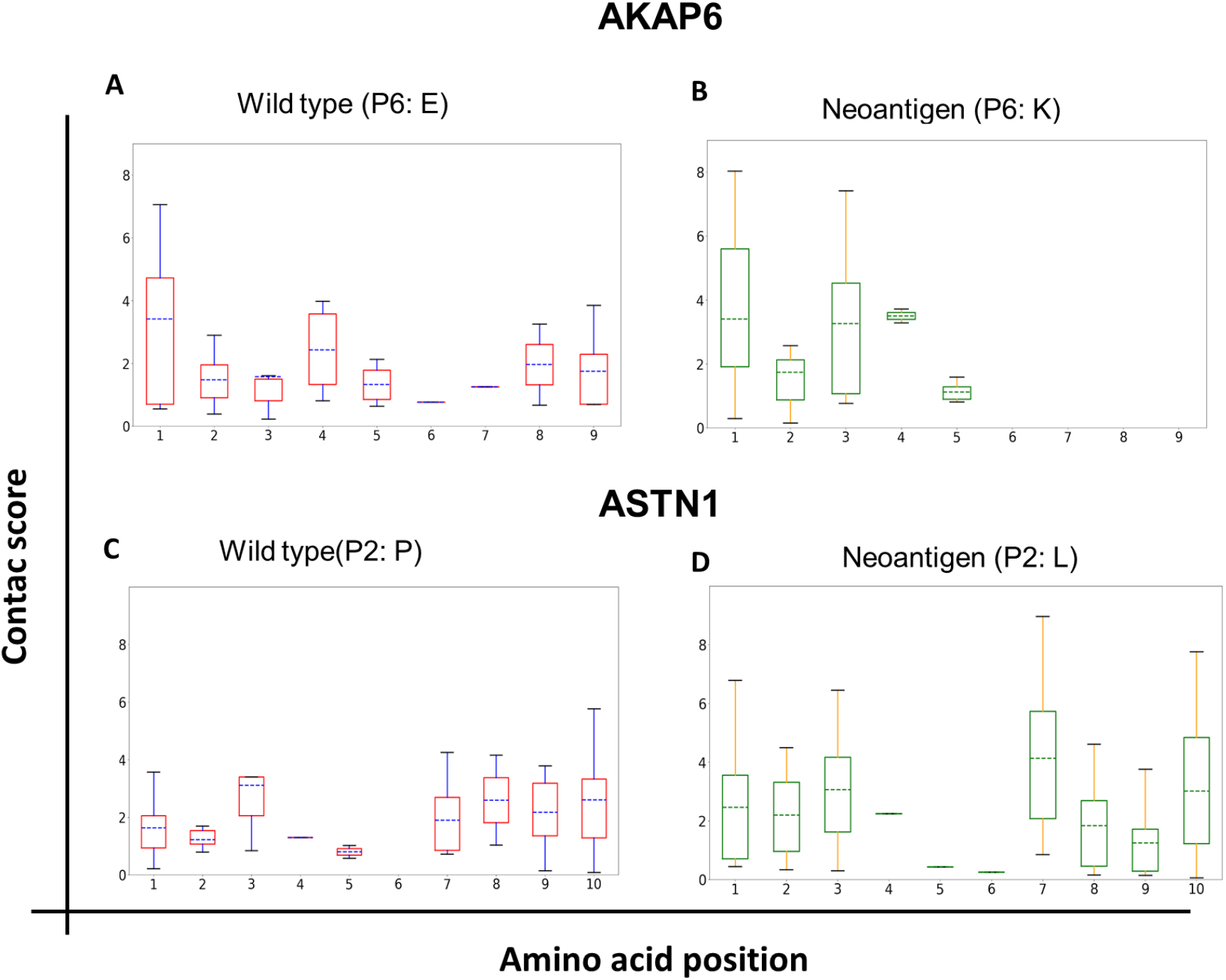
Contact scores of the interactions between amino acids of the antigens and HLA-A*02:01. Molecular dynamics simulations were performed for the wild type and the mutant version for both neoantigens, being able to analyze the intensity of the noncovalent interactions between the amino acids of the epitopes and HLA-A2. A and B box-and-whisker plots for the non-immunogenic neoantigen AKAP6. C and D box-and-whisker plots for the immunogenic neoantigen ASTN1.

As for ASTN1, the mutation in P2 generates increases in the intensity of the interaction not only at the site of the mutation but also at other positions, such as P1, P7, and P9 (see Figure 3). This result is particularly interesting because it indicates a “delocalized” (long-distance) impact of a point mutation on the global peptide-HLA interaction. In other words, a point mutation can modulate the interactions of other peptide amino acids with the PBG of the HLA molecule.

### Atomic interactions of P1 of AST1 with the HLA-A*0201 molecule

We focused the docking and molecular dynamics analyses on the atomic interactions between residues of P1 with the Peptide Binding Groove of HLA-A* 0201 in ASTN1, particularly in the ten strongest atomic interactions (see top 10 interactions in Table 4). Interactions of P1 with the HLA-A* 0201 molecule in the neoantigen were not found in the wild-type version of the sequence interacting with this molecule (whereas three out of four interactions are hydrogen bonds, the rest correspond to a different type of non-covalent interaction, see Figure 4). This observation is interesting since it allows us to measure the magnitude of the “delocalization effect” of the mutation in P2 on other positions along the peptide. By this effect, P1 becomes the position that fosters the most intense interactions stabilizing the peptide.

**Table 4:** Top 10 atomic interactions between ASTN1 and HLA-A*02:01. Highlighted in yellow are the interactions differentially expressed in the neoantigen and not in its wild-type counterpart. The nomenclature of the atoms corresponds to that used in the CHARMM36 force field.

| ASTN1 WILD TYPE |  |  |  | ASTN1 NEOANTIGEN |  |  |
| --- | --- | --- | --- | --- | --- | --- |
| # | Contact Type | Mean Score | Interaction (HLA - Peptido) | Contact Type | Mean Score | Interaction (HLA - Peptido) |
| 1 | H-bond | 0.977 | Resid. 147, atom NE1 - Resid. 9, atom O | H-bond | 0.988 | Resid. 147, atom NE1 - Resid. 9, atom O |
| 2 | Other | 0.961 | Resid. 146, atom NZ - Resid. 10, atom C | H-bond | 0.965 | Resid. 146, atom NZ - Resid. 10, atom OT2 |
| 3 | H-bond | 0.96 | Resid. 146, atom NZ - Resid. 10, atom OT1 | H-bond | 0.965 | Resid. 159, atom OH - Resid. 1, atom O |
| 4 | H-bond | 0.946 | Resid. 146, atom NZ - Resid. 10, atom OT2 | Other | 0.96 | Resid. 146, atom NZ - Resid. 10, atom C |
| 5 | Other | 0.781 | Resid. 143, atom OG1 - Resid. 10, atom C | H-bond | 0.958 | Resid. 146, atom NZ - Resid. 10, atom OT1 |
| 6 | Other | 0.778 | Resid. 146, atom CE - Resid. 10, atom OT1 | H-bond | 0.944 | Resid. 99, atom OH -Resid. 3, atom N |
| 7 | Other | 0.759 | Resid. 147, atom CD1 - Resid. 9, atom O | H-bond | 0.891 | Resid. 63, atom OE2 - Resid. 1, atom N |
| 8 | Other | 0.734 | Resid. 146, atom CE - Resid. 10, atom OT2 | H-bond | 0.881 | Resid. 63, atom OE1 - Resid. 1, atom N |
| 9 | Other | 0.715 | Resid. 147, atom NE1 - Resid. 8, atom CB | H-bond | 0.877 | Resid. 70, atom NE2 - Resid. 7, atom NE1 |
| 10 | Other | 0.694 | Resid. 146, atom CE - Resid. 10, atom C | Other | 0.871 | Resid. 63, atom CD - Resid. 1, atom N |

**Figure 4.**
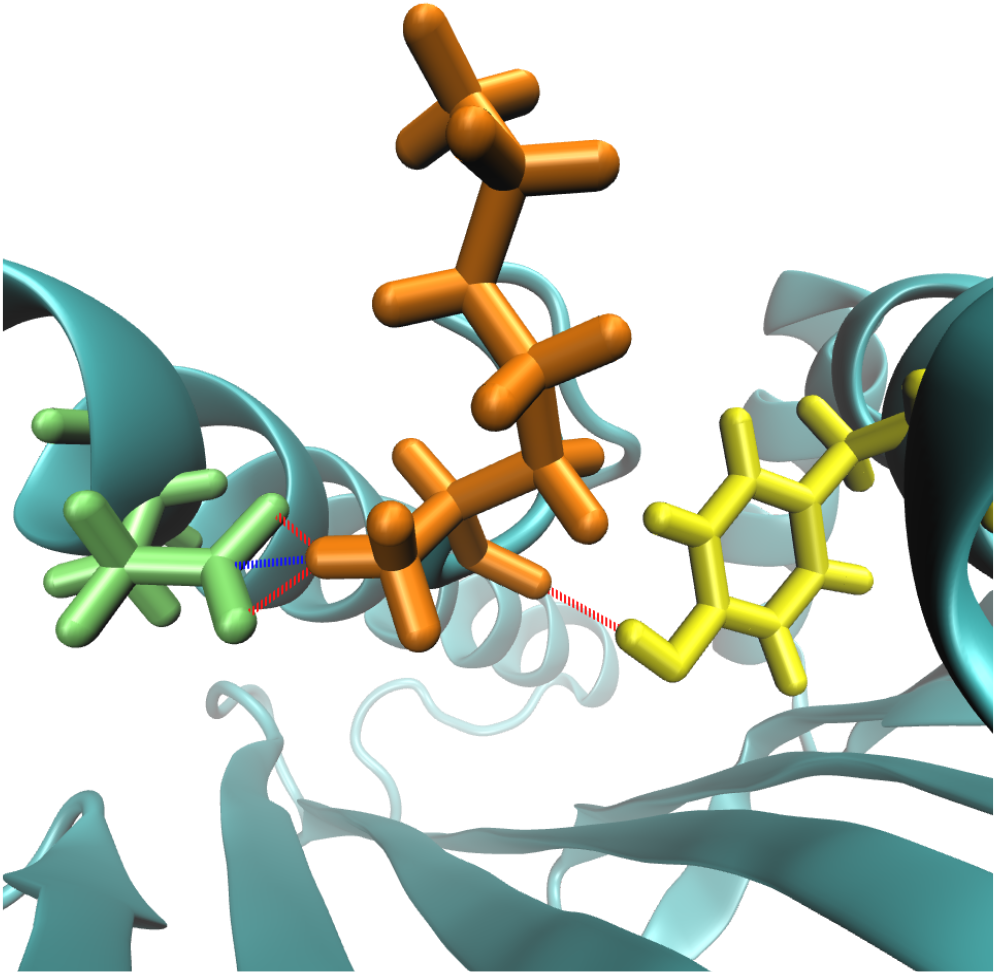
Visualization of the four interactions differentially present in the ASTN1 mutant peptide compared to its wild-type counterpart. The first amino acid of the neoantigen ASTN1 (K) is shown in orange, and the amino acids 159 and 63 of the HLA are in yellow and green, respectively. Hydrogen bonds are shown in red, and another type of interaction is shown in blue.

### Atomic interactions between P2 and its neighbors in ASTN1

As a consequence of the mutation, the intensity of the interactions of P1 with the HLA-A* 0201 changed considerably between the ASTN1 neoantigen and the wild-type sequence (Figure 3). Therefore, we investigated the effect of the mutated position on neighboring amino acids, and we found that in the wild peptide, the Proline in P2 establishes a very strong interaction with P1 that is traduced in an accumulated contact score that reaches a value of 19.14 against 13.33 in the case of the neoantigen (Supplementary Tables 2 and 3). This could explain the interactions observed at P1 of the ASTN1 neoantigen (Table 4) that are lost between the P1 of the wild-type peptide and the HLA-A* 0201 molecules, which would be hindered by proline. Regarding the interactions between P2 and P3, it is worth noting that no significant changes were found in the accumulated contact score (12.67 in the case of the wild type and 11.72 in the case of the neoantigen, see Supplementary Tables 3 and 4).

These findings highlight new features governing the complexity of peptide-HLA interactions that might explain why current bioinformatics tools are unable to accurately predict the affinity and stability of this type of complex.

### Analyses of immunogenic vs non immunogenic interactions

When comparing the non-immunogenic with the immunogenic neoantigens (AKAP6 Vs. ASTN1, respectively), it can be noticed that not only the amount and intensity of the interactions matter but also that both ends (N-terminal and C-terminal) must remain bound stably to the HLA’s PBG through the time. Also noteworthy is the fact that in the case of both AKAP6 peptides (wild type and mutant) and in the ASTN1 neoantigen, which are the peptides that remain anchored in the N-terminal part, P1 has both a significant number of interactions and high intensity (even higher than P2, Figure 3), which would indicate that this position plays a more relevant role than traditionally accepted in the stabilization of the N-terminal part of peptide epitopes.

## DISCUSSION

In the present work, we explore the use of molecular simulation techniques, such as docking and molecular dynamics, to study the interaction and stability of peptide-HLA complexes. For this, AKAP6 was selected as non-immunogenic neoantigens and ASTN1 as immunogenic according to the results obtained in healthy donors for Strønen et al.,(30) to use these tools *in-silico* and analyze the characteristics that define their immunogenicity at the pMHC complex level. When comparing the *in-vitro* immunogenicity results with the results obtained by traditional prediction tools, the failures that algorithms based solely on sequence can have been evident since AKAP6 is a clear false positive. These results can be explained by the lack of structural information on the interaction of the p-MHC complex that these tools have. Therefore, the inclusion of docking and molecular dynamics may help to strengthen the prediction of immunogenic neoantigens.

Regarding the two case selected for this work, the results provided by molecular dynamics highlight three characteristics when comparing these two neoantigens: (i) the importance of P1 and P2 for the binding of the peptide and the MHC. (ii) The delocalized effect that mutations can have and how this can influence the stability of the peptides. (iii) The importance of having high affinity and stability to the complex at both ends.

First, the results provided by the molecular dynamics of this study indicate that the P1 position of the peptide plays a key role in the stabilization of the peptide by the N-terminus than previously assumed since classically P2 and P9 were the residues of the most important peptides for anchoring to the MHC (42). This prominent role of P1 is evident in Figure 2, where significantly more important contact scores are observed in P1 than in P2 in the three peptides that remain anchored in their N-terminal part (ASTN1 neo, AKAP6 WT, and AKAP6 neo). The two neoantigens follow the classically reported amino acid sequence pattern for class I epitopes for binding to MHC-I: X-(L/I)-X(6–7)-(V/L), where L/I and V /L represent the residues whose side chain anchors the peptide to the MHC (42, 43). In the case of AKAP6, this is generated by a mutation in a non-anchor position (P6), which causes the mutant peptide to be released more quickly from the MHC through the C-terminus. On the other hand, in ASTN1, the mutation occurs right at an anchor position (P2) with a change from a Proline to a Leucine, which improves the anchoring of the peptide to the MHC and increases the number of hydrophobic interactions. These two characteristics, mutations in the anchoring residues (P2 and P9) that improve the affinity and increased hydrophobicity of the amino acids, have been reported in the literature as properties related to immunogenicity since stable interactions are generated between the anchoring amino acids and the HLA that allow a correct presentation of the antigen (44, 45).

Second, the “delocalized” impact of a point mutation on the overall peptide-HLA interaction was evidenced, meaning that a mutation can modulate the interactions of other peptide amino acids with HLA (see Table 4). This kind of effect has been previously shown in the context of interaction with the TCR since certain mutations induce structural changes in the amino acids involved in recognition by the TCR due to movement fluctuations that the side chains may have due to the mutation (46–48). That is, substitutions in a peptide can alter the intra-residual interactions that can potentially alter its conformation and, therefore, its recognition by the TCR.

Finally, the stability results of the pMHC complexes obtained agreed with the *in-vitro* immunogenicity results, which would indicate the utility of this type of in-silico strategy to identify peptides that form stable complexes with HLA proteins. When comparing the two neoantigens, it is possible to show that the immunogenic peptide (ASTN1) is linked to both ends (N-terminal and C-terminal) in a stable manner over time on HLA. These results support the importance of stability, in addition to binding affinity, as a key factor for the selection of immunogenic neoantigens (30, 33, 49), since it plays a key role in the adequate presentation of the peptide to the LT and, thus trigger an immune response. This is reinforced by previous studies that have reported failures in the selection of neoantigens when the main or only parameter considered is the binding affinity (23, 50–52).

Even though only two peptides were analyzed, the results shed light on characteristics of immunogenicity; however, these must be validated in larger cohorts to define the role they have in binding affinity and stability in the MHC, and finally, on immunogenicity. However, due to the considerable computational cost of this type of strategy, its use would be restricted to the final stages of the immunogenic neoantigen identification pipelines, where a small number of candidates remain. These tools have the potential to work not only at the level of the interaction within the peptide and the MCH but also to determine the interactions between the p-MHC complex and TCR, which can eventually be implemented to select the best TCRs for adoptive therapy purposes.

## CONCLUSIONS

Personalized cancer vaccines are presented as a novel and promising alternative to cancer immunotherapy, especially in those cases where effective treatments do not yet exist. Currently, most selection of neoantigens is made by *in silico* methodologies (15), however, the results of clinical studies reveal that the *in vivo* immunogenicity of *in silico* epitopes is very limited. Therefore, it is important to search for new tools that allow the selection of immunogenic epitopes that yield better clinical results *in vivo* when used as a vaccine. An important attribute of immunogenic MHC-peptide complexes is the long half-life these complexes have (33, 49, 53–55), so we believe that molecular simulation can play an important role in the fine-tuning of the selection process required to select neoantigens to be included in a neoantigen vaccine. In the present work, we explored the use of molecular simulation techniques such as docking and molecular dynamics to analyze the role of peptide-HLA complex stability in immunogenicity for CD8+ T lymphocytes. The analysis of the stability of the analyzed complexes collated with the results of immunogenicity *in vitro* leads to confirm that kind of relationship. These results point to the suitability of this type of *in silico* strategy to identify peptides that form stable complexes with HLA proteins that are highly immunogenic for CD8+ T cells.

Regarding the case study selected for this work, in addition to relating the stability of HLA-peptide complexes with immunogenicity, the results provided by molecular dynamics indicate that the P1 position of the peptide plays a more important role in the stabilization of the N-terminal part of what was assumed until now. Likewise, the results suggest that the mutations may have a “delocalized effect” on the peptide-HLA interaction; that is, they may modulate the intensity of the interactions of distant amino acids of the peptide throughout the PBG of the HLA molecules.

## ACKNOWLEDGMENT

This study was supported by funding from the Universidad Nacional de Colombia. DIB, Vicedecanatura de Investigación Universidad Nacional Medical School; funds from two joint grants among Fundación Salud de Los Andes, Universidad Nacional, and COLCIENCIAS (see funding section below. Experiments presented in this paper were carried out using the Grid’5000 testbed, supported by a scientific interest group hosted by Inria and including CNRS, RENATER and several Universities as well as other organizations (see https://www.grid5000.fr). The authors express their gratitude to “Laboratorio en Investigación en Sistemas inteligentes – LISI” and the leader of this group Luis Fernando Niño, “Grupo de Investigación en Bioinformática y Biología de Sistemas GiBBS” and their leader Andrés Pinzón and Janeth Gonzales for the advice in the analysis of the molecular docking results.

## DATA AVAILABILITY

Docking and molecular dynamics simulation data of AKAP6 and ASTN1 peptides are available at Zenodo with https://doi.org/10.5281/zenodo.5772726.

## SUPPLEMENTARY MATERIAL

### Access to molecular dynamics simulations: https://doi.org/10.5281/zenodo.5772726

**Table S1.**
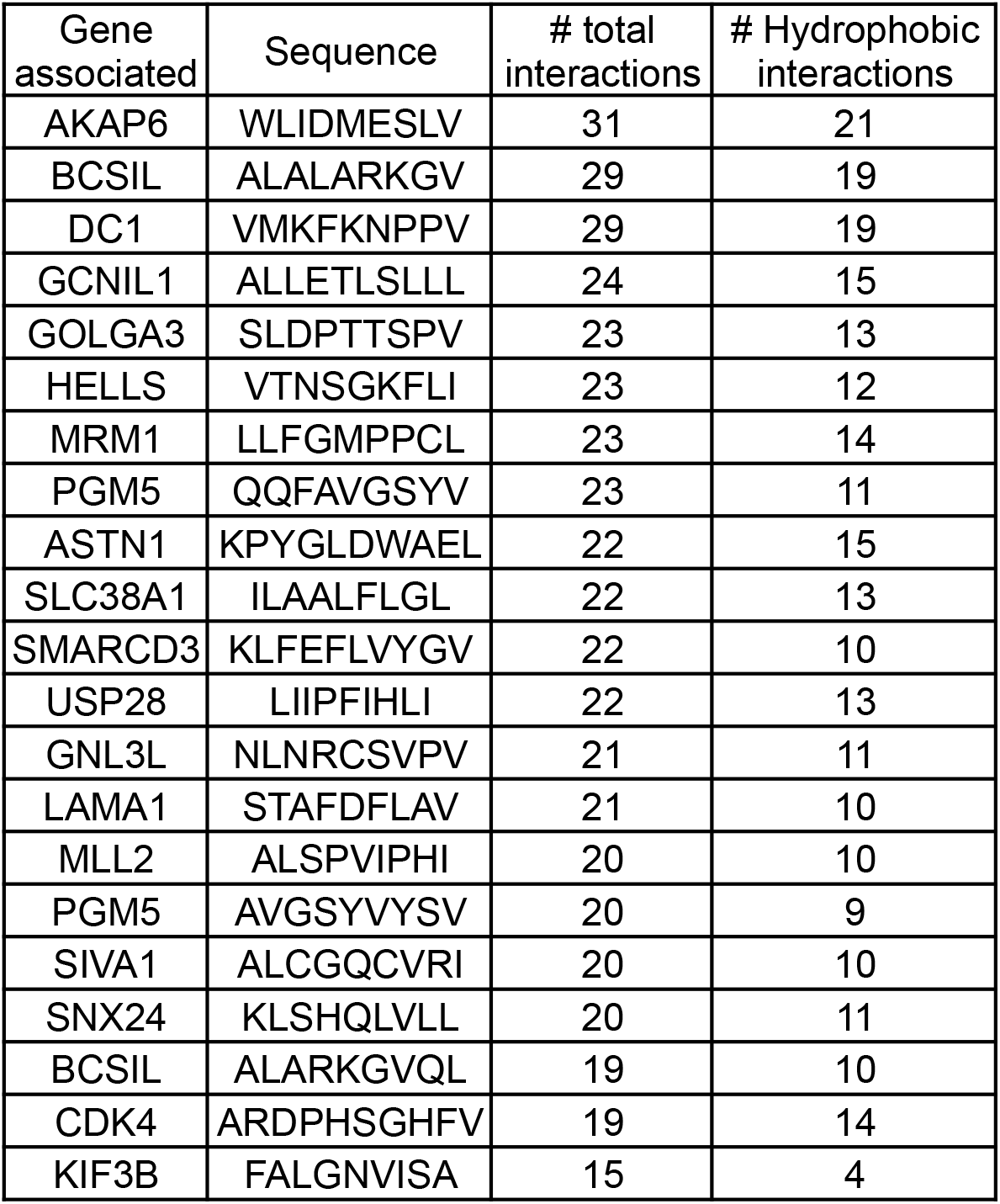
Number of interactions in between the peptide and HLA.

**Table S2:** Interactions between P1 and P2 in the ASTN1 wild type

| # | P1 | P2 | Contact score |
| --- | --- | --- | --- |
|  | Atom | Atom |  |
| 1 | C | N | 1 |
| 2 | O | N | 1 |
| 3 | CA | N | 1 |
| 4 | C | CA | 1 |
| 5 | O | CA | 0,997 |
| 6 | C | C | 0,979 |
| 7 | C | O | 0,95 |
| 8 | O | C | 0,946 |
| 9 | N | N | 0,943 |
| 10 | CB | N | 0,846 |
| 11 | O | O | 0,807 |
| 12 | C | CB | 0,78 |
| 13 | CA | CA | 0,698 |
| 14 | CG | N | 0,34 |
| 15 | O | CB | 0,257 |
| 16 | C | CG | 0,251 |
| 17 | CA | O | 0,228 |
| 18 | CA | C | 0,099 |
| 19 | CB | O | 0,08 |
| 20 | O | CG | 0,056 |
| 21 | N | CA | 0,034 |
| 22 | CB | CA | 0,024 |
| 23 | CA | CB | 0,012 |
| **Accumulated contact score** |  |  | **13,33** |

**Table S3:** Interactions between P2 and P3 in the ASTN1 neoantigen

| # | P2 | P3 | Contact score |
| --- | --- | --- | --- |
|  | Atom | Atom |  |
| 1 | C | N | 1 |
| 2 | O | N | 1 |
| 3 | CA | N | 1 |
| 4 | C | CA | 0,999 |
| 5 | O | CA | 0,997 |
| 6 | C | C | 0,983 |
| 7 | CB | N | 0,982 |
| 8 | O | C | 0,975 |
| 9 | C | O | 0,895 |
| 10 | N | N | 0,834 |
| 11 | O | O | 0,812 |
| 12 | C | CB | 0,771 |
| 13 | CA | CA | 0,685 |
| 14 | O | CB | 0,219 |
| 15 | CB | O | 0,144 |
| 16 | CG | N | 0,124 |
| 17 | CB | CA | 0,1 |
| 18 | CA | O | 0,073 |
| 19 | CA | C | 0,072 |
| 20 | CB | C | 0,026 |
| 21 | CA | CB | 0,013 |
| 22 | N | CA | 0,011 |
| **Accumulated contact score** |  |  | **11,72** |

**Table S4:** Interactions between P1 and P2 in the ASTN1 neoantigen

| # | P1 | P2 | Contact score |
| --- | --- | --- | --- |
|  | Atom | Atom |  |
| 1 | C | N | 1 |
| 2 | O | N | 1 |
| 3 | C | CA | 0,999 |
| 4 | C | CD | 0,999 |
| 5 | CA | N | 0,999 |
| 6 | O | CA | 0,997 |
| 7 | CA | CD | 0,992 |
| 8 | C | C | 0,99 |
| 9 | O | C | 0,985 |
| 10 | C | O | 0,962 |
| 11 | O | O | 0,925 |
| 12 | O | CD | 0,895 |
| 13 | N | N | 0,887 |
| 14 | CB | N | 0,87 |
| 15 | C | CG | 0,852 |
| 16 | C | CB | 0,841 |
| 17 | CA | CA | 0,678 |
| 18 | N | CD | 0,581 |
| 19 | CG | O | 0,496 |
| 20 | CA | O | 0,416 |
| 21 | CB | CD | 0,403 |
| 22 | CG | N | 0,393 |
| 23 | CB | O | 0,275 |
| 24 | O | CB | 0,242 |
| 25 | CA | C | 0,18 |
| 26 | CA | CG | 0,101 |
| 27 | O | CG | 0,081 |
| 28 | CB | CA | 0,041 |
| 29 | CB | C | 0,031 |
| 30 | CA | CB | 0,019 |
| 31 | N | CA | 0,014 |
| **Accumulated contact score** |  |  | **19,14** |

**Table S5:** Interactions between P2 and P3 in the ASTN1 wild type

| # | P2 | P3 | Contact score |
| --- | --- | --- | --- |
|  | Atom | Atom |  |
| 1 | C | N | 1 |
| 2 | O | N | 1 |
| 3 | C | CA | 0,999 |
| 4 | CA | N | 0,999 |
| 5 | O | CA | 0,997 |
| 6 | C | C | 0,977 |
| 7 | O | C | 0,953 |
| 8 | CB | N | 0,95 |
| 9 | N | N | 0,836 |
| 10 | C | CB | 0,826 |
| 11 | O | O | 0,716 |
| 12 | C | O | 0,712 |
| 13 | CA | CA | 0,677 |
| 14 | O | CB | 0,464 |
| 15 | C | CG | 0,333 |
| 16 | CA | C | 0,059 |
| 17 | CG | N | 0,055 |
| 18 | CB | CA | 0,046 |
| 19 | CD | N | 0,029 |
| 20 | CA | CB | 0,027 |
| 21 | N | CA | 0,012 |
| **Accumulated contact score** |  |  | **12,67** |

